# Patterns of transcriptional response to 1,25-dihydroxyvitamin D3 and bacterial lipopolysaccharide in primary human monocytes

**DOI:** 10.1101/030759

**Authors:** Silvia N. Kariuki, John D. Blischak, Shigeki Nakagome, David B. Witonsky, Anna Di Rienzo

**Affiliations:** Department of Human Genetics, University of Chicago, Chicago, IL, USA 60637

**Author notes:** **Correspondence:** Prof. Anna Di Rienzo, Department of Human Genetics, The University of Chicago, 920 E. 58th Street, Room 424, Chicago, IL 60637, USA.

**Keywords:** differential expression profiling, innate immune cells, pro-inflammatory response, pathway analysis

## Abstract

The active form of vitamin D, 1,25-dihydroxyvitamin D3 (1,25D), plays an important immunomodulatory role, regulating transcription of genes in the innate and adaptive immune system. The present study examines patterns of transcriptome-wide response to 1,25D and the bacterial lipopolysaccharide (LPS) in primary human monocytes, to elucidate pathways underlying the effects of 1,25D on the immune system. Monocytes obtained from healthy individuals of African-African and European-American ancestry were treated with 1,25D alone or in the presence of LPS, which induced significant up-regulation of genes in the antimicrobial and autophagy pathways, while pro-inflammatory response genes were significantly down-regulated. A joint Bayesian analysis enabled clustering of genes into patterns of shared transcriptional response across treatments. The biological pathways enriched within these expression patterns highlighted several mechanisms through which 1,25D could exert its immunomodulatory role. Pathways such as mTOR signaling, EIF2 signaling, IL-8 signaling and Tec Kinase signaling were enriched among genes with opposite transcriptional responses to 1,25D and LPS, respectively, highlighting the important roles of these pathways in mediating the immunomodulatory activity of 1,25D. Furthermore, a subset of genes with evidence of inter-ethnic differences in transcriptional response was also identified, suggesting that in addition to the well-established inter-ethnic variation in circulating levels of vitamin D, the intensity of transcriptional response to 1,25D and LPS also varies between ethnic groups. We propose that dysregulation of the pathways identified in this study could contribute to immune-mediated disease risk.

## Introduction

Vitamin D plays an important immunomodulatory role through a transcriptional mechanism (1–4). In monocytes/macrophages, activation of toll-like receptors (TLRs) by bacterial ligands such as lipopolysaccharide (LPS) induces *CYP27B1,* which encodes the enzyme that converts inactive 25-hydroxyvitamin D3 (25D) to active 1,25-dihydroxyvitamin D3 (1,25D) (1, 5) inside the cell. 1,25D binds the vitamin D receptor (VDR) and translocates into the nucleus, modulating transcription of vitamin D-responsive genes that are involved in various immune pathways (3, 4).

In monocytes/macrophages, 1,25D induces antimicrobial peptide genes such as cathelicidin antimicrobial peptide (*CAMP*) (1, 2, 5), defensin genes such as β-defensin 4A (*DEFB4A*) (6), and autophagy genes such as autophagy related 5 (*ATG5*) (1, 7). Several lines of epidemiological evidence suggest that intracrine production of 1,25D from circulating inactive 25D in monocytes/macrophages is important for this direct antimicrobial activity. Insufficient circulating levels of 25D have been linked to increased susceptibility to tuberculosis (Tb) (8, 9). Supplementation with vitamin D3 in individuals with insufficient levels of serum 25D resulted in an enhanced antimicrobial response (1, 5, 10).

Although many studies have been conducted on the inter-individual and inter-ethnic variation in the circulating inactive 25D levels, relatively little is known about variation in the transcriptional response to active 1,25D. Treatment of THP-1 macrophage cell lines with 1,25D enhanced the cytokine/chemokine response to *Mycobacterium tuberculosis* (*M. tb*) infection (11), while 1,25D treatment of peripheral blood mononuclear cells (PBMCs) downregulated the expression of genes in immune-related pathways such as interferon signaling (12). In dendritic cells (13, 14), 1,25D treatment reduced their ability to promote T cell activation, thereby attenuating the pro-inflammatory response (14, 15). The mechanism underlying this tolerogenic profile involved metabolic reprogramming towards oxidative phosphorylation (14). The immunoregulatory role of 1,25D in different innate immune cell types is complex, but it generally involves the attenuation of an intense pro-inflammatory response, which can have toxic consequences such as sepsis and septic shock (16-18).

The present study focuses on the transcriptome-wide response to 1,25D and LPS in primary human monocytes *in vitro,* enabling characterization of the transcriptional response to 1,25D in the context of a pro-inflammatory stimulus. We identify biological pathways enriched among genes with opposite transcriptional responses to 1,25D and LPS, such as EIF2 signaling and mTOR signaling, which highlight some mechanisms through which 1,25D could exert its immunomodulatory role. In addition, we identify some inter-ethnic differences in response patterns, suggesting that the well-established inter-ethnic variations in the vitamin D pathway extend to the intensity of transcriptional response to LPS and 1,25D.

## Materials and Methods

### Ethics Statement

All donors to Research Blood Components (http://researchbloodcomponents.com/) and Sanguine Biosciences (https://www.sanguinebio.com/) sign an IRB-approved consent form giving permission to collect blood, and use it for research purposes. This study did not require IRB review at the University of Chicago because blood samples were not shipped with individually identifiable information.

### Subjects

All subjects were healthy donors collected by Research Blood Components and Sanguine Biosciences. Self-reported ethnicity, age, gender, date, and time of blood drawing were recorded for each donor. Buffy coats from 10 AA and 10 EA subjects were shipped within 24 hours of collection. We processed samples in multiple batches, balanced by ethnic group. Serum 25D levels and parathyroid hormone (PTH) levels were determined at the Clinical Chemistry Laboratory of the University of Chicago using standard assays (cat. no. 06506780160 and cat. no. 11972103160 respectively, Roche Diagnostics Corporation, Indianapolis, IN, USA).

### Monocyte culture and treatment

We isolated peripheral blood mononuclear cells (PBMCs) from the buffy coats of the 20 subjects by density gradient centrifugation using Ficoll-Paque PLUS medium (GE Healthcare Life Sciences, Pittsburgh, PA). We isolated monocytes from the PBMCs by positive selection using magnetic CD14 MicroBeads according to the supplier’s protocol (Miltenyi Biotec, San Diego, CA). We cultured isolated monocytes (1×10^6^ cells/mL) in RPMI 1640 medium (Gibco, Life Technologies, Grand Island, NY), 25mg/mL Gentamicin (Gibco) and 10% charcoal-stripped fetal bovine serum (Gibco) in 24-well plates. Monocytes were cultured in three replicates for 24 hours for each of the following treatments: 1) Vehicle solution containing 1% Ethanol and 99% culture medium, as a negative control (E), 2) 100nM of 1,25D, 3) 10ng/mL of LPS in the vehicle solution, and 4) 100nM of 1,25D and 10ng/mL of LPS (1,25D+LPS) (experimental design summarized in **Fig. S1**).

### Transcriptome analysis

We pooled replicates for each treatment and extracted total RNA from the pool using Qiagen RNeasy Plus mini kit (Valencia, CA). We extracted RNA from 80 samples consisting of the 20 subjects that each received 4 treatments, in 10 batches each balanced by ethnic group. RNA concentration and RNA integrity score (RIN) were recorded for each sample on the 2100 Bioanalyzer instrument (Agilent Technologies, Santa Clara, CA) (average RNA concentration and RIN scores in each ethnic group summarized in **Table S1**). Total RNA was reverse transcribed into cDNA, labeled, hybridized to Illumina (San Diego, CA, USA) Human HT-12 v3 Expression Beadchips and scanned at the University of Chicago Functional Genomics Core facility. The microarrays were hybridized in three batches, and we recorded the array batch number each sample, to be used as a covariate in subsequent analyses. We performed low-level microarray analyses using the Bioconductor software package lumi (19) in R, as previously described (20). Briefly, we annotated probes by mapping their sequence to RefSeq (GRCh37) transcripts using BLAT. We discarded probes that mapped to multiple genes to avoid ambiguity in the source of a signal due to cross-hybridization of similar RNA molecules. We also discarded probes containing one or more HapMap SNPs to avoid spurious associations between expression measurements and ethnicity, due to allele frequency differences between ethnic groups. We applied variance stabilization to all arrays, discarded poor quality probes, and quantile normalized the arrays using the default method implemented in the lumiN function. After these filters, probes mapping to 10,958 genes were used in downstream analyses.

### Differential expression analysis

We tested each gene for differential expression (DE) using a linear mixed-effects model with the R package, lme4 (21). The model included fixed effects for ancestry and the three treatment conditions (1,25D, LPS, 1,25D+LPS), as well as interaction effects between ancestry and the treatments. It also included a random effect to model the differences between the individuals. Lastly, the model included covariates for the technical factors with the strongest effects on the expression data (p < 0.05), as determined by their association with the principal components described below, including array batch, age, baseline 25D levels, baseline PTH levels, RNA concentration and RIN scores. P-values were obtained using the R package, lmerTest, which provides a summary function with p-values added for the t-test based on the Satterthwaite approximation for denominator degrees of freedom (22). To correct for multiple testing, we estimated the false discovery rate (FDR) using the “qvalue” function in R, based on the Storey method (23). The FDR for DE was set at 1%. To identify genes that were DE between the two ancestries, we tested the significance of the interaction effects. Here we used a more relaxed FDR threshold of 10% to determine significance, due to the smaller sample size in the inter-ethnic comparison (10 AA’s and 10 EA’s).

We also performed a joint Bayesian analysis using the R package Cormotif (24), which jointly models expression data across different experiments enabling classification of genes into patterns of shared and distinct differential expression. Genes are assigned to correlation motifs, which are the main patterns of differential expression obtained from the shared information across experiments, which in our study are treatments and ethnic groups. We regressed out the technical covariates described above from the expression data using the limma package removeBatchEffect (25), and used the residuals as input. We used a modified version of Cormotif as described in (26) where the original code was modified to return the cluster likelihood for each gene to enable downstream analyses. Also, since Cormotif is non-deterministic, we ran each test 100 times and kept the result with the largest maximum likelihood estimate.

### Gene set enrichment analysis

We performed gene set enrichment analyses using the commercially available software Ingenuity Pathway Analysis (IPA). We compared DE genes with curated functional attribution lists organized by canonical pathway function. The magnitude of over-representation of a particular canonical pathway in the gene list from our study was calculated as the ratio of the number of genes from our data set that map to the pathway divided by the total number of reference genes in that pathway in the IPA database. Statistical significance of the observed enrichment of a particular pathway was determined using Benjamini-Hochberg multiple testing corrected p-values provided by IPA (27).

### Identifying vitamin D receptor binding sites near DE genes

We reanalyzed published data sets of VDR ChIP-seq, which used THP-1 monocytic cell lines treated with 1,25D and LPS or 1,25D alone (33), and FAIRE-seq, which used THP-1 cells treated with 1,25D (34). First, we aligned sequence reads to the human reference (GRCh37) using BWA backtrack 0.7.5 (28). Second, we kept only sequence reads with phred-scaled mapping quality ≥ 30 using samtools v1.1 (29). Third, PCR duplicate were removed with picard tool v 1.130 (http://broadinstitute.github.io/picard/). For the ChIP-seq data sets, we confirmed the quality of data sets by strand cross-correlation (SCC) analysis (30) implemented in the R script “run_spp_nodups.R” packaged in phantompeakqualtools (https://code.google.com/p/phantompeakqualtools/). Statistically significant peaks were identified using MACS version 2 (31) with the following essential command line arguments: macs2 callpeak --bw X -g hs --qvalue=0.05 -m 5 50, where X is a length of the bandwidth that was defined as a fragment length calculated by SCC for the ChIP-seq data or as 200 bp for the FAIRE-seq data reported in Seuter *et al.* (2012).

To identify VDR response elements, we considered peaks that overlapped completely or partially between the ChIP-seq data after 1,25D and LPS treatment and the FAIRE-seq data. We then annotated them using HOMER (32) to find the closest gene to each peak and, among these genes, we selected those that were DE genes in response to the combined 1,25D+LPS treatment from the linear mixed-effects analysis. Enrichment of VDR response elements was determined using Fisher’s exact test, comparing peaks in DE genes to those in non-DE genes.

We also examined the enrichment of VDR binding sites among genes in different Cormotif expression patterns from the joint Bayesian analysis. To inclusively identify VDR binding sites, we merged the ChIP-seq data after 1,25D and LPS treatment and data from cells treated with 1,25D alone. We examined overlap between the genes that were closest to the peaks, and the genes in each Cormotif pattern. Enrichment of VDR peaks was then determined using Fisher’s exact test, comparing VDR peaks in each Cormotif pattern with peaks in the “Non-DE” Cormotif pattern.

## Results

### Sources of transcriptome-wide variation

We performed principal components analysis (PCA) of the variance-stabilized log_2_-transformed expression data using the prcomp function in R. Principal component 1 (PC1) separates the samples by LPS treatment, accounting for 22% of the total variation in gene expression and reflecting the large effect of LPS on the transcriptome. (**Fig. S2 (A) and (C), Table S2**) and PC2 separates the samples by 1,25D treatment, and accounts for 8.6% of the total variation in gene expression (**Fig. S2 (A) and (D), Table S2**). PC3 and PC4, which account for 6.7% and 5.8% of variation respectively, separate the samples by the three array processing batches (**Fig. S2 (B), (E) and (F), Table S2**). We also tested for associations between the PCs and the different covariates recorded for each sample (average sample covariates are compared between ancestries in **Table S1**). PC1 was associated with RNA concentration (p = 1.54E-05, r^2^ = 0.219), PC2 was weakly associated with the RNA integrity number (RIN) scores (p = 0.009, r^2^ = 0.086), while PC3 was associated with age (p = 1.36E-06, r^2^ = 0.266), baseline 25D levels (p = 0.001, r^2^ = 0.136), baseline PTH levels (p = 6.1E-06, r^2^ = 0.237), and RIN score (p = 0.005, r^2^ = 0.099) (**Table S2**). The effects of array processing batch, RNA concentration, RIN scores, serum 25D and PTH levels were subsequently included as covariates in the linear mixed-effects analysis for differential expression.

After regressing out the covariates using the limma package removeBatchEffect, and performing PCA on the residuals of the covariates-corrected expression data, we observed that PC1 and PC2 separated the samples by treatment (**Fig. S3 (A), (C) and (D), Table S3**), but PC1 was still associated with RNA concentration (p = 3.56E-05, r^2^ = 0.203) while PC2 was still associated with RIN score (p = 0.003, r^2^ = 0.113) (**Table S3**). PC3, which accounted for 5% of the total variation in gene expression, was associated with sample (p = 4.5E-07, r^2^ = 0.286), and ancestry (p = 1.04E-05, r^2^ = 0.227), highlighting the effect of inter-individual and inter-ethnic variation on gene expression (**Fig. S4, Table S3**).

### Opposite effects of 1,25D and LPS on the transcriptome

Using the main effects for each treatment from the linear mixed-effects model, we identified genes that were DE in response to the different treatments at a FDR of 1%. 2,888 genes were DE in response to 1,25D alone relative to vehicle. Gene set enrichment analysis identified metabolic processes, such as oxidative phosphorylation and the tricarboxylic acid (TCA) cycle, enriched among up-regulated genes (**Table S4**). Pathways that play important roles in regulating translation processes, such as EIF2 signaling and mTOR signaling, were also significantly enriched among up-regulated genes, indicating an important role of 1,25D in regulating translation. Immune responses involving chemokine signaling, B and T cell signaling, as well as various pro-inflammatory signaling cascades such as Tec kinase signaling, Phospholipase C signaling, and Integrin signaling were enriched amongst the down-regulated genes, consistent with the immunomodulatory function of 1,25D.

There was a strong transcriptomic response to LPS treatment relative to vehicle, with 4,461 genes DE at a FDR of 1%. Pathways enriched among LPS responsive genes highlight the opposite direction of transcriptional response to 1,25D and LPS, where pro-inflammatory immune response pathways were enriched among up-regulated genes, while oxidative phosphorylation and translational control pathways were enriched among down-regulated genes (**Table S4**), indicating the importance of these pathways in the pro-inflammatory effects induced by LPS stimulation.

### Effects of combined 1,25D+LPS treatment on the transcriptome

The combined treatment of 1,25D+LPS resulted in 4,720 genes significantly DE relative to vehicle. We also examined the transcriptional response of the combined 1,25D+LPS treatment relative to LPS in an attempt to isolate the effect of 1,25D on the transcriptome in the presence of LPS. 2,404 genes were significantly DE in the combined 1,25D+LPS relative to LPS.

The pattern of response to 1,25D+LPS treatment relative to vehicle followed a similar pattern to LPS treatment alone, probably because of the overwhelming effect of LPS on transcriptional response in monocytes. Similar pathways were also enriched among genes in these treatment categories (**Table S4**). Similarly, when we analyzed the response to 1,25D+LPS treatment relative to LPS, which effectively subtracts the transcriptional effects of LPS, the pathways enriched among DE genes were similar to those for 1,25D treatment relative to vehicle.

Additional pathways were enriched among genes responsive to the combined 1,25D+LPS treatments, both relative to vehicle and relative to LPS; these included adipogenesis and insulin receptor signaling pathways, both involved in lipid metabolic processes, which were enriched among up-regulated genes. IL-4 signaling, which is associated with allergy and asthma through development of T cell mediated immune responses (33, 34), was significantly enriched among down-regulated genes (**Table S4**). Pathways enriched among genes responsive to the combined 1,25D+LPS treatment indicate a regulatory role of 1,25D in these pathways specifically in the context of LPS stimulation.

### Bayesian analysis of shared transcriptional response across treatments and ethnic groups

To further dissect the effects of 1,25D and LPS on the transcriptome, we sought to identify the shared and distinct patterns of transcriptional response across treatments and across ethnic groups. A popular approach to this question is to investigate the overlap of DE genes between conditions at a given FDR threshold. However, this approach fails to account for incomplete power to detect DE genes, thus exaggerating the differences in the transcriptional response between the conditions. In order to identify shared patterns of transcriptional response across treatments and ancestry while accounting for incomplete power, we implemented a joint Bayesian analysis with the R/Bioconductor package Cormotif (24). Genes were classified into different response patterns, or Cormotifs, across treatments and ancestry (**Fig. 1**). Since Cormotif does not distinguish the direction of effect across treatments, we used the results of the linear mixed-effects analysis in conjunction with the Cormotif approach to establish direction of response in the different patterns (**Fig. 2**).

**Fig. 1:**
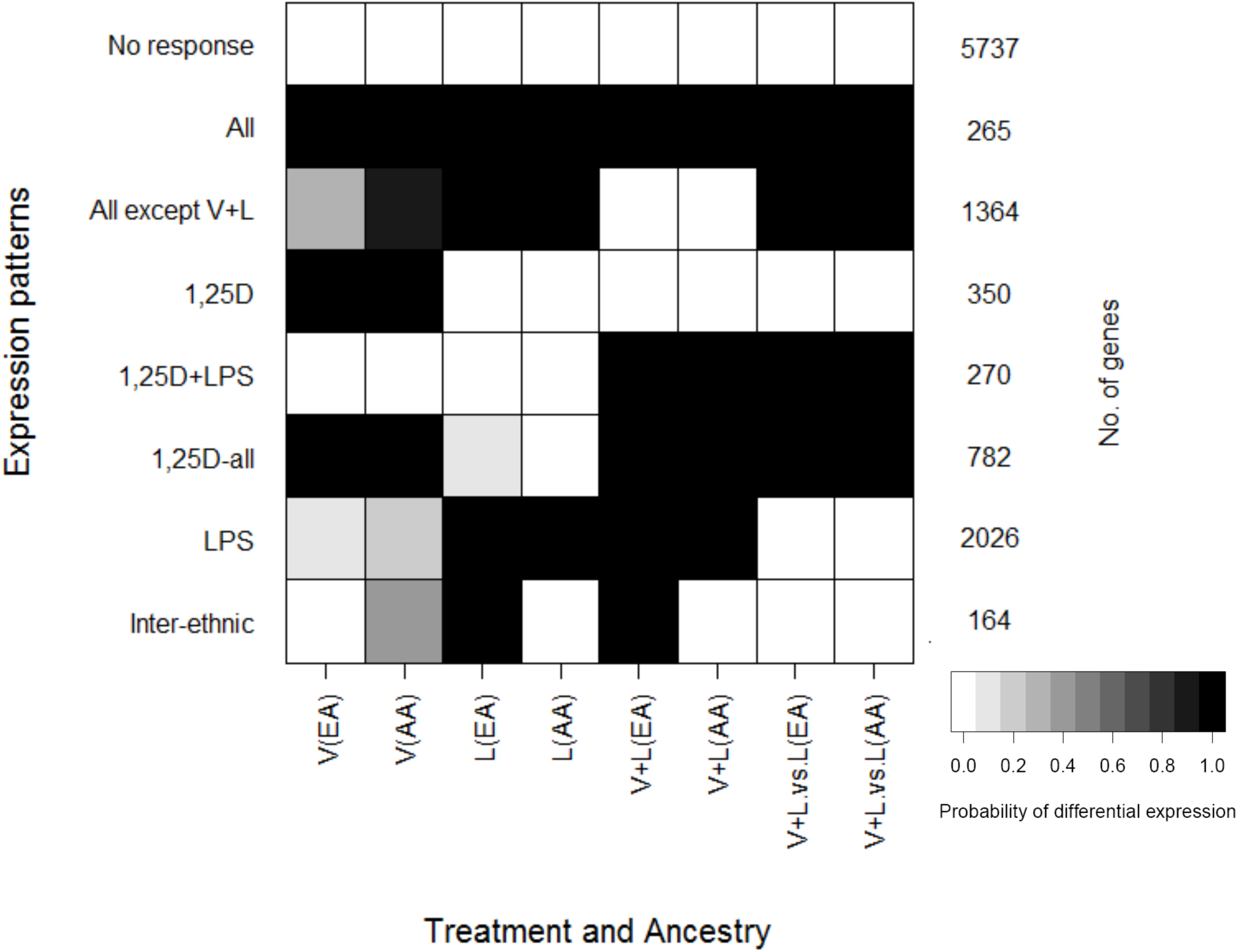
Joint Bayesian analysis identifies transcriptional response patterns across the different treatments and ancestries. The shading of each box represents the posterior probability that a gene assigned to a given expression pattern (rows) is differentially expressed in individuals from a particular ancestry in response to each treatment (columns). **V**=Response to 1,25D, relative to vehicle; **L** = Response to LPS relative to vehicle; **V+L** = Response to 1,25D+LPS relative to vehicle; **V+L.vs.L** = Response to 1,25D+LPS relative to LPS; **EA** = European-American; **AA** = African-American

**Fig. 2:**
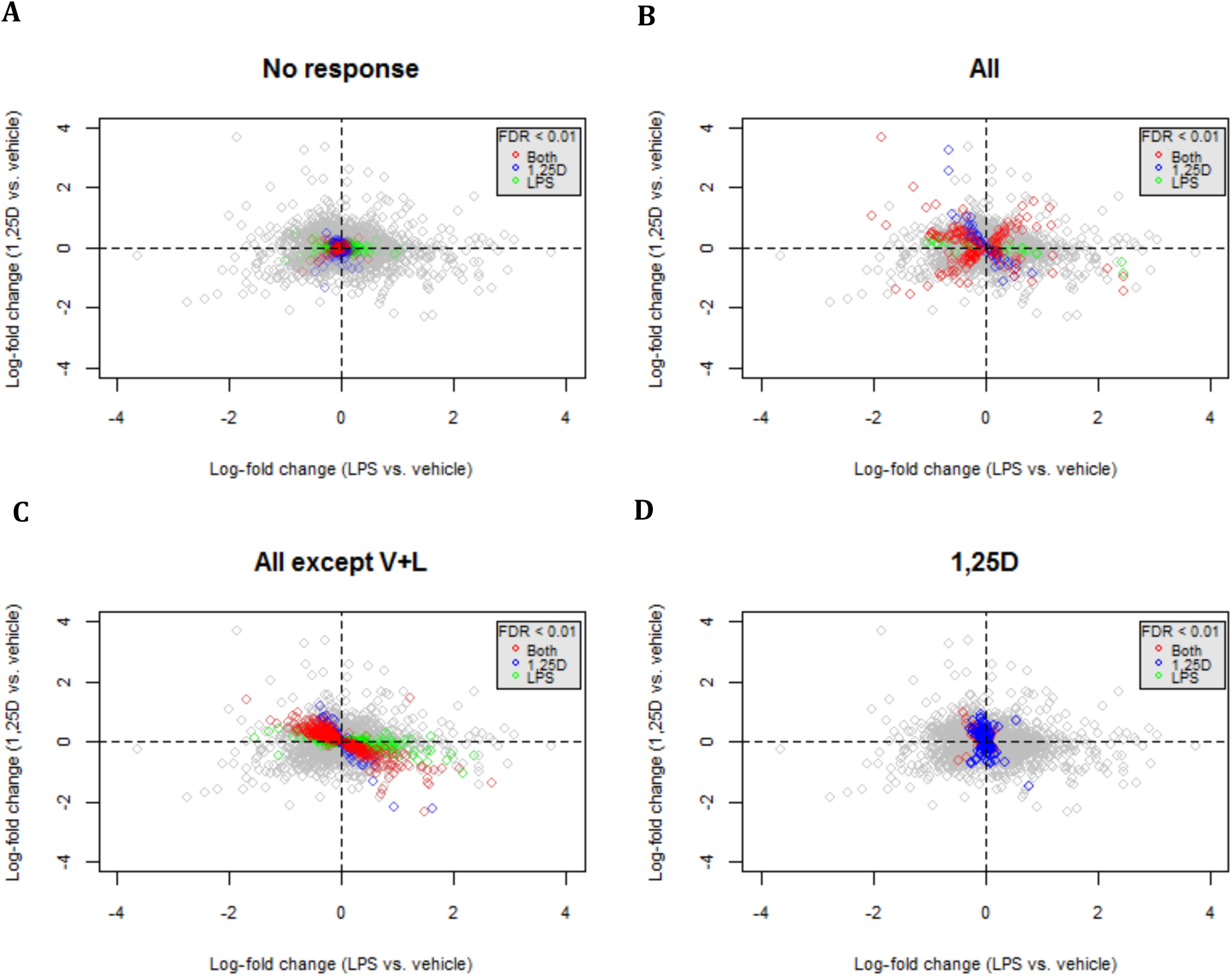
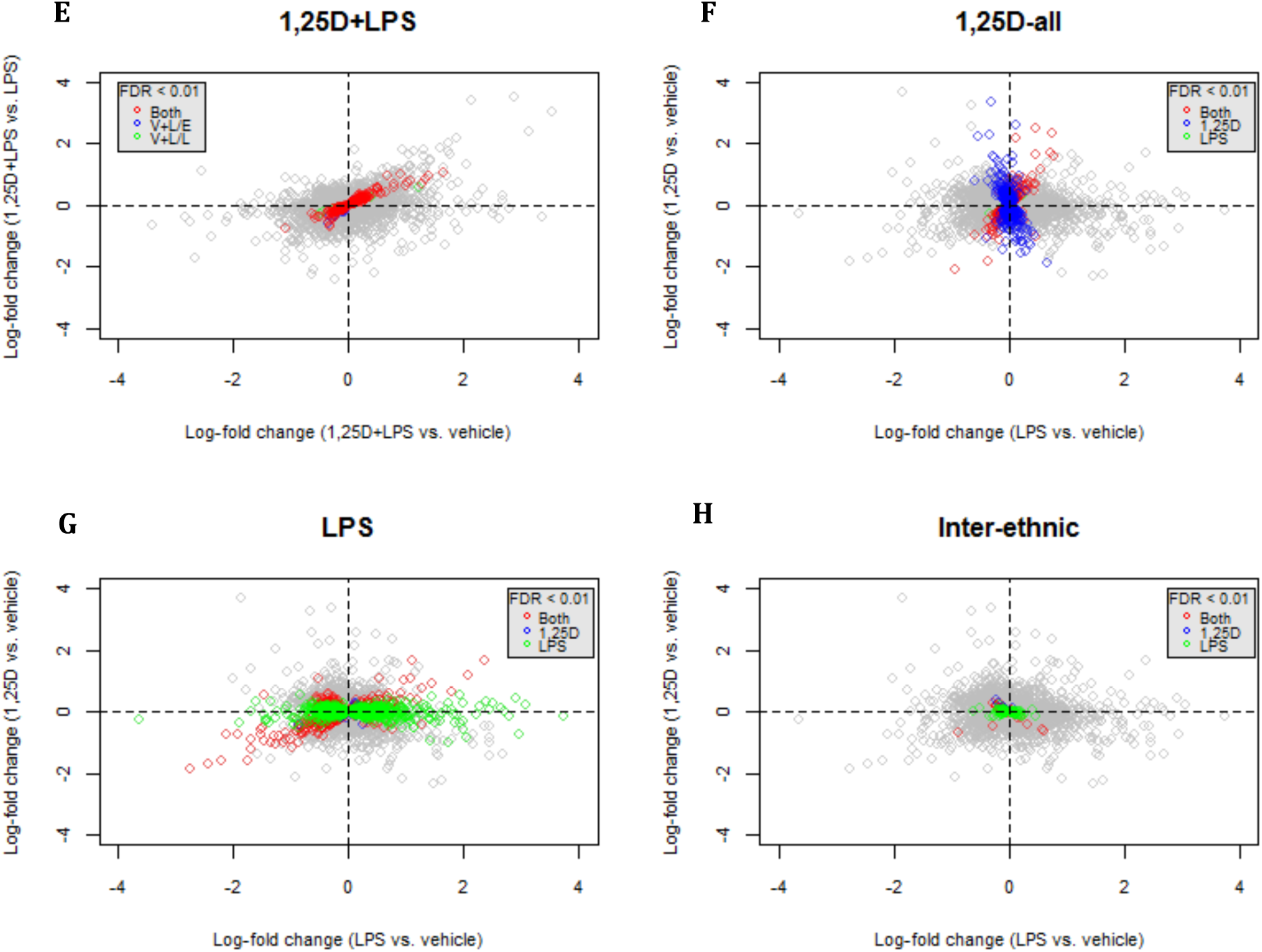
Patterns of differential response to single treatment with 1,25D (vertical axis) or LPS (horizontal axis) for each Cormotif shown in A-D and F-H. E shows patterns of differential response to the combined treatment with 1,25D and LPS relative to LPS (vertical axis) and 1,25D and LPS relative to vehicle (horizontal axis). Genes are color coded based on q-values < 0.01 from linear mixed-effects analysis as follows: **Red** = DE in response to both 1,25D and LPS; **Blue** = DE in response to 1,25D; **Green** = DE in response to LPS; **Grey** = not DE.

A total of 5,737 genes were classified in the “No response” pattern, which includes genes whose expression levels were unchanged across all the treatments (**Figs. 1 and 2A**). This is broadly consistent with the results of the linear mixed-effects analysis, with 80% of these genes being also classified as non-DE for any treatment in the linear mixed-effects analysis.

Genes that responded to all the treatments were classified in the “All” pattern and included 265 genes whose expression levels changed across all treatments and ancestries (**Figs. 1 and 2B**). Genes classified in this Cormotif had response patterns to 1,25D and LPS that were both concordant (i.e. up- or down-regulated in both treatments) and discordant (i.e. up-regulated in one treatment and down-regulated in the other). Genes that were up-regulated in all treatments (top-right quadrant, **Fig. 2B**) included *CD14,* which encodes a surface antigen expressed on monocytes that is involved in mediating response to bacterial LPS. Genes that were down-regulated in all treatments included chemokine signaling genes such as *CCL13* (bottom-left quadrant, **Fig. 2B**). The discordant response patterns included genes that were up-regulated by 1,25D and down-regulated by LPS (top-left quadrant, **Fig. 2B**), with EIF2 signaling and mTOR signaling pathways significantly enriched among these genes (**Table 1**). This is consistent with the opposite transcriptional effects of 1,25D and LPS on genes in these pathways that were highlighted in the linear mixed-effects analysis. Genes that were down-regulated by 1,25D and up-regulated by LPS (bottom-right quadrant, **Fig. 2B**) included some cytokine receptor genes such as *IL7R* and *IL2RA* which are important components of the pro-inflammatory signaling cascade.

**Table 1:**
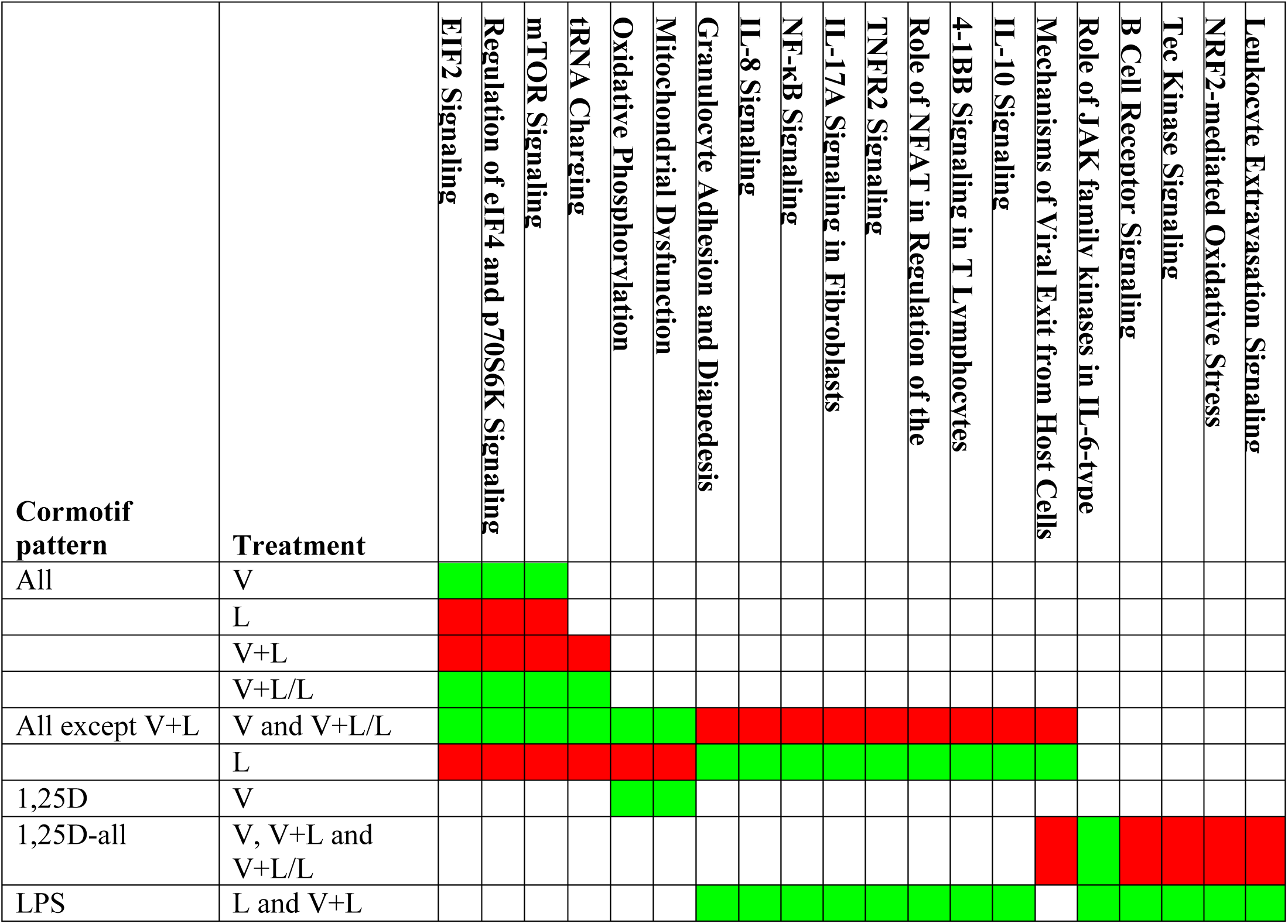
Sharing of enriched biological pathways across Cormotifs. The table shows the biological pathways that are enriched (FDR < 0.05) in more than one Cormotif subdivided based on the direction of transcriptional response (up-regulated genes in **green** and down-regulated genes in **red**) and the treatment (**V**=Response to 1,25D, relative to vehicle; **L** = Response to LPS relative to vehicle; **V+L** = Response to 1,25D+LPS relative to vehicle; **V+L/L** = Response to 1,25D+LPS relative to LPS).

The “All except V+L” pattern included 1,364 genes whose expression levels changed in all treatments except the combined 1,25D+LPS relative to vehicle. All the genes in this Cormotif pattern were discordant in their response to 1,25D and LPS resulting in a neutral effect in the response to the combined 1,25D+LPS treatment relative to vehicle (**Figs. 1 and 2C**). Genes that were responsive to the combined 1,25D+LPS treatment relative to LPS followed a similar direction of response to the individual 1,25D response genes, suggesting that the response to 1,25D at these genes is not dramatically influenced by LPS. The genes that were up-regulated by 1,25D and down-regulated by LPS (top-left quadrant, **Fig. 2C**) were enriched for EIF2 and mTOR signaling pathways, similar to the discordant genes in the “All” category (**Table 1**). In addition, oxidative phosphorylation and mitochondrial dysfunction pathways were significantly enriched amongst these genes. On the other hand, genes that were down-regulated by 1,25D and up-regulated by LPS (bottom-right quadrant, **Fig. 2C**) were enriched for various pro-inflammatory response pathways, including Granulocyte Adhesion and Diapedesis, IL-8 signaling, NF-kB signaling, TNFR2 signaling and Role of NFAT in regulation of the immune response (**Table 1**).

Genes responsive to 1,25D were divided into three Cormotif patterns: “1,25D”, “1,25D+LPS”, and “1,25D all”. The “1,25D” pattern included 350 genes that were DE in response to 1,25D alone, in the absence of LPS (**Figs. 1 and 2D**). Oxidative phosphorylation and mitochondrial dysfunction pathways were significantly enriched among up-regulated genes (**Table 1**). Interestingly, the oxidative phosphorylation pathway genes enriched in this Cormotif pattern responded similarly to the genes in the same pathway classified in the “All except V+L” Cormotif pattern, in that they are significantly induced by 1,25D. However, the oxidative phosphorylation pathway genes in the “1,25D” pattern respond exclusively to 1,25D, while those in the “All except V+L” Cormotif pattern are up-regulated by 1,25D and down-regulated by LPS (**Table 1, Fig. S5**). This indicates a context-specific response profile among genes in the same pathway, where some genes in the oxidative phosphorylation pathway are uniquely regulated by 1,25D, whereas other genes in the same pathway are regulated by both 1,25D and LPS.

The “1,25D+LPS” pattern included 270 genes that responded to 1,25D only in the presence of LPS. This pattern captured genes that were DE in response to the combined 1,25D+LPS treatment relative to both vehicle and to LPS (**Figs. 1 and 2E**). Although there were no enriched pathways among genes in this Cormotif pattern at an FDR of 5%, some interesting pathways, such as eNOS signaling and cholesterol biosynthesis pathway, were represented among the down-regulated genes at a FDR of 27%, suggesting a role for 1,25D in modulating these pathways upon LPS stimulation.

The “1,25D all” pattern included 782 genes that were DE in response to 1,25D in the presence and in the absence of LPS (**Figs. 1 and 2F**). Genes in the antimicrobial pathway were included in this category, such as the anti-bacterial peptide gene *CAMP,* autophagy genes *ATG3, ATG5, ATG2A* and *ATG9A* and the intracellular pattern recognition receptor gene *NOD2.* These genes were significantly up-regulated in response to 1,25D alone or in combination with LPS. The Role of JAK family kinases in IL-6-type Cytokine Signaling was the most significantly enriched pathway amongst the up-regulated genes (**Table 1**), and included molecules such as *MAPK14, PTPN11* and *STAT5B,* all of which could be crucial for triggering antimicrobial responses in monocytes. Biological pathways enriched among down-regulated genes in this category included B cell receptor signaling, Tec kinase signaling and Leukocyte extravasation signaling (**Table 1** and **Fig. S6**), highlighting the role of 1,25D in repressing pro-inflammatory response pathways. Interestingly, immunological and inflammatory diseases were among the most enriched disease categories among the down-regulated genes (**Table S6** and **Fig. S6**), suggesting a direct protective role of 1,25D in immunological diseases. Overall, the “1,25D all” response pattern illustrates the important dual immunomodulatory role played by 1,25D in monocytes, where antimicrobial pathway genes are up-regulated, while pro-inflammatory pathway genes associated with immunological and inflammatory disease are down-regulated by 1,25D in the presence or absence of LPS stimulation.

The “LPS” pattern included 1,400 genes whose expression levels changed in response to LPS treatment and the combined 1,25D+LPS treatment relative to vehicle (**Figs. 1 and 2G**). Consistent with the results from the linear mixed-effects analysis, pro-inflammatory pathways were significantly enriched among the up-regulated genes in this category, including IL-8 signaling, NF-kB signaling, IL-17 signaling, and TNFR2 signaling among others (**Table 1**). Among the down-regulated genes, tRNA charging, mitochondrial dysfunction, the TCA Cycle II, galactose metabolism pathway and folate transformation pathway were significantly enriched (**Table S5**), indicating that LPS modulates transcription of genes in these metabolic pathways.

### Genes with inter-ethnic differential response

The “Inter-ethnic” pattern was of particular interest, as it identified 164 genes with evidence of differential responses to LPS and the combined 1,25D+LPS treatment relative to vehicle between AA’s and EA’s, with a stronger response in EA’s compared to AA’s (**Figs. 1 and 2H**). Though no biological pathway was enriched among these genes at a FDR < 5%, some interesting immune response pathways were represented among the up-regulated genes such as IL-10 signaling and TNFR2 signaling (at a FDR of 14%), suggesting that the pro-inflammatory effects of LPS are stronger in EA’s than in AA’s.

We also interrogated the degree of inter-ethnic differences in transcriptional response using the main interaction term for treatment and ancestry in the linear mixed-effects analysis. We identified 16 genes with strong inter-ethnic differences in response to the combined 1,25D+LPS treatment relative to vehicle at a FDR < 10%. These genes include *STEAP3,* which encodes an endosomal ferrireductase required for efficient transferrin-dependent iron uptake, *PPAP2B* which encodes a member of the phosphatidic acid phosphatase (PAP) family and has been implicated in coronary artery disease risk (35, 36), and *AKNA* which encodes a transcription factor that specifically activates the expression of the CD40 receptor and its ligand CD40L/CD154 on lymphocyte cell surfaces, which are critical for antigen-dependent-B-cell development (**Fig. 3A**). Interestingly, 13 out of the 16 genes showed more significant differential responses in EA’s (**Fig. S7**), similar to the pattern observed in the “Inter-ethnic” Cormotif.

**Fig. 3:**
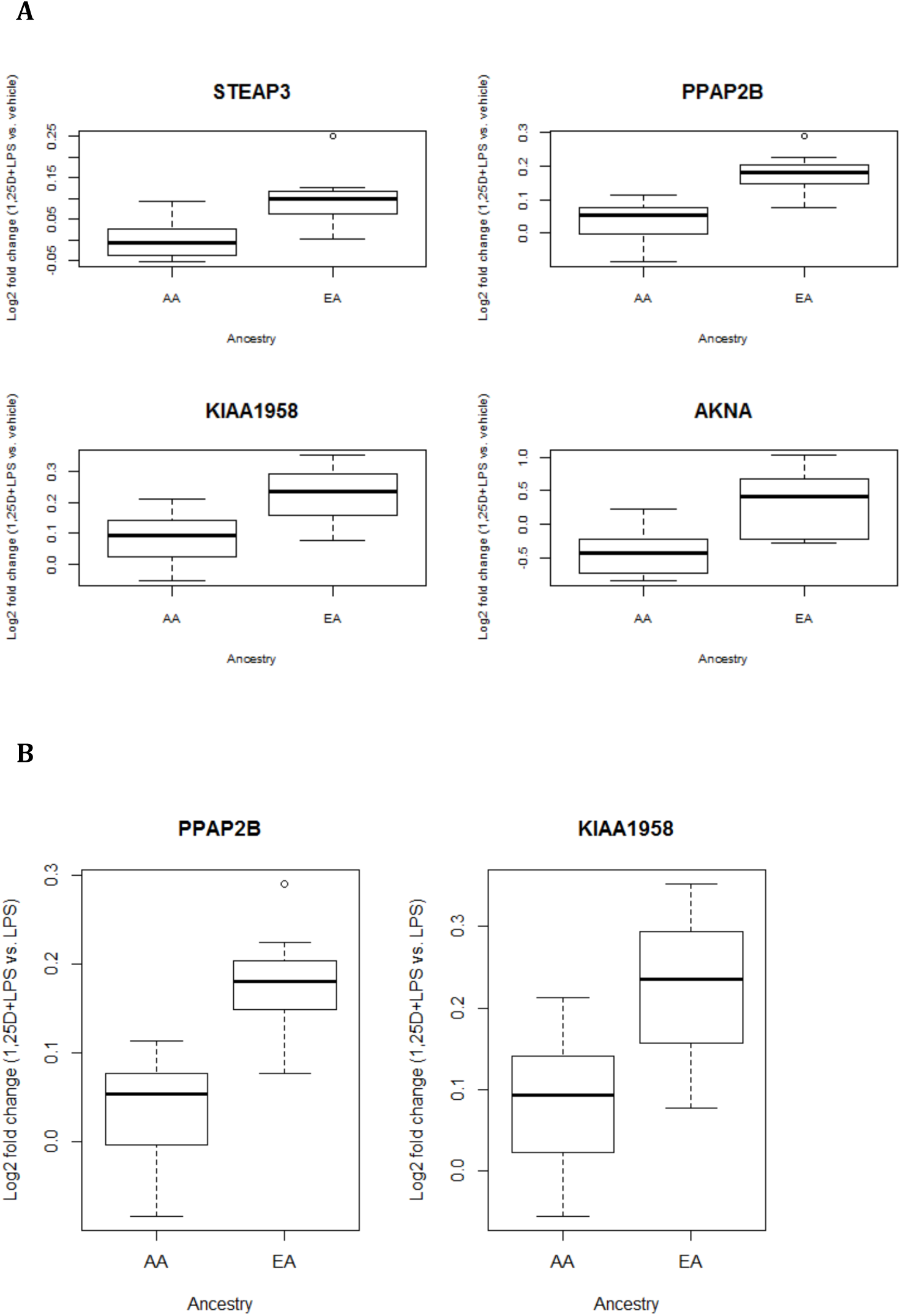
Genes with inter-ethnic differences in transcriptional responses to 1,25D+LPS relative to vehicle (**A**), and relative to LPS (**B**) were identified using the interaction term for treatment and ancestry in the linear mixed-effects model (FDR < 0.10). The boxplots show examples of these genes with different log fold change in transcript levels between the two ethnic groups. **AA** = African-American; **EA** = European-American.

To account for the effect of LPS, we also examined inter-ethnic differential response to the combined 1,25D+LPS treatment relative to LPS. *PPAP2B* and *KIAA1958* were the only statistically significant genes identified in this category (p = 9.66E-06 and 1.13E-05, respectively), and again these two genes were more significantly differentially expressed in EA’s (**Fig. 3B**).

### Regulatory elements near DE genes

We examined the overlap between genes DE in response to the combined 1,25D+LPS treatment in our primary monocytes and published datasets for VDR ChIP-seq (37) and FAIRE-seq (38) performed in THP-1 cells to examine whether there was enrichment of open chromatin regions and VDR binding sites near the transcription start sites of DE genes. We found a significant enrichment of VDR binding sites amongst genes DE in response to the combined 1,25D+LPS treatment relative to vehicle (p = 3.97E-08) and relative to LPS (p = 3.19E-11) (**Table S7**). There was an overlap of 201 genes between the three datasets, highlighting genes such as *CAMP* and *CD14,* which contain open chromatin regions and VDR binding sites near the transcription start site under regulation by 1,25D; these 201 genes are potentially direct VDR targets.

In addition, we examined the enrichment of VDR binding sites across the different Cormotif patterns (**Table 2**). The genes in the “1,25D-all” and “All” Cormotif patterns had the highest enrichment of VDR binding sites (p = 6.88E-13 and 2.57E-08 respectively), indicating a higher proportion of potentially direct VDR targets represented in these Cormotif patterns. Interestingly, genes in the “1,25D” Cormotif pattern were not significantly enriched for VDR binding sites, suggesting that the presence of LPS, in addition to 1,25D, is important to enable the 1,25D-VDR transcriptional activity in primary monocytes.

**Table 2:** Proportion of genes in each Cormotif pattern containing VDR binding sites. Enrichment of VDR peaks in each category calculated using Fisher’s exact test, comparing genes in each Cormotif pattern to those in the “Non-DE” Cormotif pattern.

| **Cormotif pattern** | **Total No. Genes** | **No. Genes with VDR binding site** | **Proportion of genes with VDR binding site** | **Enrichment p-value** |
| --- | --- | --- | --- | --- |
| Non-DE | 5737 | 186 | 0.03 | - |
| All | 265 | 31 | 0.12 | 2.57E-08 |
| All except V+L | 1364 | 65 | 0.05 | 0.01 |
| 1,25D | 350 | 17 | 0.05 | 0.13 |
| 1,25D+LPS | 270 | 23 | 0.09 | 1.49E-04 |
| 1,25D-all | 782 | 76 | 0.10 | 6.88E-13 |
| LPS | 2026 | 96 | 0.05 | 3.79E-03 |
| Inter-ethnic | 164 | 8 | 0.05 | 0.27 |

## Discussion

We used a transcriptomic approach to characterize the immunomodulatory role of 1,25D in the presence of a pro-inflammatory stimulus to identify the mechanisms through which 1,25D exerts its immunomodulatory role. We analyzed differential expression patterns using both a linear mixed-effects analysis modeling individual treatment comparisons, and a Bayesian analysis using the Cormotif method, which jointly models differential expression across all treatments and ethnic groups, thereby accounting for incomplete power. A similar joint Bayesian framework has been successfully applied to expression quantitative trait loci (eQTL) mapping to distinguish between shared and context-specific eQTLs (39, 40). Our joint Bayesian analysis enabled clustering of DE genes into distinct transcriptional response patterns, with pathways enriched within these transcriptional patterns highlighting mechanisms that mediate the immunomodulatory role of 1,25D.

Metabolic pathways involving oxidative phosphorylation were enriched among up-regulated genes in the “All except V+L” and “1,25D” Cormotif patterns (**Table 1**). We highlight context-specific response pattern of genes within this pathway, where some genes were uniquely induced by 1,25D, while genes in other parts of the pathway were regulated by both 1,25D and LPS. The crucial role played by 1,25D in regulating oxidative phosphorylation was previously reported in PBMCs and dendritic cells (12-14), and this regulation of metabolic reprogramming by 1,25D is thought to be crucial for controlling function, growth, proliferation, and survival of various immune cell subsets (41, 42). The fact that LPS down-regulated genes in the oxidative phosphorylation pathway confirms previous reports indicating that LPS induces a metabolic shift away from oxidative phosphorylation to anaerobic glycolysis in macrophages and dendritic cells to enable ATP production (42). This effect is similar to the Warbug effect in tumor cells whose high energy demand is met by switching the metabolic profile away from the tricarboxylic acid cycle and the oxidative phosphorylation pathway, towards glycolysis thereby enabling rapid ATP production (42, 43). Previous work done in mouse macrophages and dendritic cells (44, 45) indicated that a metabolic shift towards glycolysis mediated the pro-inflammatory response, and this pro-inflammatory response could be attenuated by pharmacologic inhibition of glycolysis. From our study, this subset of oxidative phosphorylation pathway genes that were down-regulated by LPS, were then up-regulated by addition of 1,25D in combination with LPS (**Table 1**). Therefore, oxidative phosphorylation could be one of the mechanisms through which 1,25D attenuates the pro-inflammatory response induced by LPS in monocytes, and the subset of genes in this pathway that we identified which were modulated by both LPS and 1,25D could be central to this mechanism.

The mTOR signaling pathway was consistently enriched among genes that were up-regulated by 1,25D and down-regulated by LPS in the “All” and “All except V+L” patterns. mTOR signaling was previously implicated in inhibition of pro-inflammatory response in LPS-stimulated monocytes/macrophages and dendritic cells, as well as in the maintenance of a tolerogenic phenotype in dendritic cells (14, 46-48). Inhibition of mTOR resulted in increased pro-inflammatory cytokine production by LPS-stimulated monocytes/macrophages and dendritic cells (46, 48) and increased T cell proliferation (14, 48), implicating a role of mTOR in regulating the pro-inflammatory response. The genes in this pathway were significantly down-regulated by LPS; however this direction of response was reversed by addition of 1,25D in combination with LPS (**Table 1**), implying that 1,25D attenuates the pro-inflammatory response by up-regulating mTOR signaling. In addition, the genes in this pathway play important roles in regulating translation initiation, and include the ribosomal protein gene *RPS27,* and the eukaryotic translation initiation factor gene *EIF2A,* which encodes the eukaryotic initiation factor 2 (eIF-2α) that has been shown to be a downstream target of the vitamin D receptor (49). Therefore, regulation of translation initiation through targeting the mTOR signaling pathway could be a novel mechanism for the attenuation of the pro-inflammatory response mediated by 1,25D in monocytes.

Furthermore, the EIF2 signaling pathway was also enriched among genes up-regulated by 1,25D and down-regulated by LPS in the “All” and “All except V+L” patterns, and this result is consistent with the individual treatment DE analysis using the linear mixed-effects model (**Tables 1, S4 and S5**). EIF2 signaling plays an important role in regulating translation initiation in response to stress, and was implicated in regulating pro-inflammatory cytokine production and bacterial invasion in mouse embryonic fibroblast cells (MEFs) (50). Shrestha *et al.* (2012) reported that the Yersinia-encoded virulence factor, YopJ, inhibited EIF2 signaling in MEFs. Similarly in our study, LPS consistently down-regulated genes in the EIF2 signaling pathway, in a mechanism that might be similar to that triggered by YopJ. In addition, Shrestha *et al.* (2012) observed that mutant MEFs with defective EIF2 signaling that were infected with different bacterial pathogens experienced enhanced cytotoxicity compared to wild type, due to increased bacterial invasion, indicating a direct role of EIF2 signaling in the antimicrobial response. 1,25D could hence exert its antimicrobial role in monocytes by up-regulating genes in the EIF2 signaling pathway.

The dual immunomodulatory role of 1,25D was also highlighted by the genes clustered in the “1,25D-all” pattern. While 1,25D broadly down-regulated genes in the pro-inflammatory cytokine and signaling cascade pathways, and indeed had a protective role against various immunological diseases that were associated with down-regulated genes in this category, it also played a crucial role in inducing important antimicrobial and autophagy genes in this Cormotif pattern. The most significantly enriched biological pathway among the up-regulated genes was the Role of JAK family kinases in IL-6-type cytokine signaling, which contained genes such as *STAT5B* which regulates signaling in diverse biological processes. Previous reports indicate that the TLR2/1-mediated induction of the vitamin D-dependent antimicrobial pathway requires IL-15 activity (51), which could be mediated via STAT5 activation which has been shown to be important for IL-15 signaling (52, 53). 1,25D could hence regulate genes in this pathway to trigger antimicrobial responses in monocytes.

By profiling transcriptional response in monocytes from individuals of African-American and European-American ancestries, we identified some patterns of inter-ethnic variation in response to LPS, and the combined 1,25D+LPS treatment in both the linear mixed-effects analysis and the joint Bayesian analysis, while correcting for inter-individual variation in baseline levels of circulating 25D. This raises the intriguing possibility that inter-ethnic variation in the vitamin D pathway is not limited to the well-established differences in circulating levels of 25D (54-56), but it may extend to the intensity of the transcriptional response to LPS and 1,25D. Interestingly, most of the genes with inter-ethnic differential response showed more significant differential responses in EA’s. The fact that most of the inter-ethnic transcriptional differences were detected in the response to LPS or to the combined 1,25D+LPS, both relative to vehicle, suggests that these two ethnic groups differ in the pro-inflammatory response. However, two genes had significant inter-ethnic differences in transcriptional response to the combined 1,25D+LPS relative to LPS (**Fig. 3B**), suggesting that they differ more specifically in their response to vitamin D.

We further identified enrichment of VDR ChIP-seq and FAIRE-seq peaks among genes DE in response to the combined 1,25D+LPS treatments. This enrichment was particularly strong for genes in the “1,25D-all” and “All” Cormotif patterns, suggesting that a substantial proportion of these genes are under direct regulation of the 1,25D-VDR transcription factor complex. It further highlights the importance of a combination of both 1,25D and LPS in stimulating the transcriptional activity of the 1,25D-VDR transcriptional complex in primary human monocytes. It is possible that a larger fraction of VDR binding sites could be found near DE genes if we assessed the transcriptional response in different conditions. Specifically, while we cultured primary monocytes with 1,25D and LPS for 24 hours, the VDR ChIP-seq data was collected in THP-1 monocytic cell lines cultured with 1,25D and LPS for about 2 hours (37). Future VDR ChIP-seq studies in primary monocytes will enable better characterization of the regulatory architecture of 1,25D response genes, and more accurate conclusions about direct and indirect targets of the 1,25D-VDR transcription factor complex.

Overall, through transcriptomic profiling, our study characterizes the dual immunomodulatory role of 1,25D in primary human monocytes, highlighting the importance of biological pathways such as mTOR signaling and EIF2 signaling in mediating this immunomodulatory role. The pathways highlighted in this study may provide mechanistic clues for the observed associations between insufficient levels of circulating serum 25D and increased disease risk. The inter-individual and inter-ethnic variation in intracellular transcriptional response to 1,25D has not been previously characterized, and could serve as an additional contribution to disease risk.

## Acknowledgements

The authors are grateful to Carole Ober, Marcelo Nobrega, and John Novembre, for advice on analysis methods, and Sonia Kupfer, Choongwon Jeong and the entire Di Rienzo lab for helpful discussions about the project. This work was supported in part by NIH grant (R01 GM101682) to AD. SNK was supported by the Howard Hughes Medical Institute Gilliam Fellowship for Advanced Study. This project was supported by the University of Chicago Comprehensive Cancer Center Support Grant (#P30 CA14599), with particular support from the Genomics Core Facility.

